# Impaired chromosome segregation results in sperms with excess centrosomes in *emb-27^APC6^* mutant *C. elegans*

**DOI:** 10.1101/449538

**Authors:** Tomo Kondo, Akatsuki Kimura

## Abstract

The anaphase-promoting complex (APC) is a major regulator of chromosome segregation and is implicated in centriole engagement, whose de-regulation causes abnormal number of centrosomes. The *emb-27* gene in *C. elegans* encodes a subunit of APC. The paternal *emb-27* mutant was reported to show cell division with multiple furrows, suggesting the presence of excess centrosomes. In this study, we examined the number of centrosomes and the mechanism underlying de-regulation of centrosome number in *emb-27* mutants. Our observations indicated excess centrosomes in *emb-27* sperms, which resulted in zygotes with excess centrosomes. Further, the secondary spermatocyte of *emb-27* produced reduced number of spermatids, which is likely the direct cause of the excess number of centrosomes per sperm. We propose that a chromosome segregation defect in *emb-27* induced centrosome separation defect, resulting in reduced number of buds. Additionally, treatment with cytoskeletal inhibiting drugs indicated presence of three kinds of forces working in combination to move the centrosomes into the spermatids. The present study suggested a novel role of microtubule in the budding cytokinesis of spermatocytes.

## Introduction

The centrosome is a major microtubule-organizing center in animal cells (Stearns and Winey, 1997). Normally dividing cells have 2 centrosomes, each containing a pair of centrioles, that become the 2 poles of the bipolar mitotic spindle to segregate the sister chromatids into 2 daughter cells after mitosis (Mazia, 1987). The number of centrosomes in a cell is strictly regulated (Hinchcliffe and Sluder, 2001; Nigg and Holland, 2018), and an excess number of centrosomes or centrioles is known to be deleterious to cells (Pihan, 2013). Each pair of centrioles duplicate only once during a cell cycle (Azimzadeh and Bornens, 2007), similar to chromosomes that are also duplicated only once during a cell cycle, which is critical for cell proliferation and for preventing aneuploidy (Storchova and Kuffer, 2008). Interestingly, centrioles and chromosomes use common molecules—cohesin and its regulators—to connect pairs of sister centrioles/chromosomes together and to release the connection at specific timing during cell cycle. The anaphase-promoting complex (APC) plays a role in separation of the sister chromatids in anaphase by degrading securin, which induces activation of separase and subsequently the cleavage of cohesin to facilitate the separation of sister chromatids (Kamenz and Hauf, 2017; Uhlmann et al., 1999; Yeong, 2004). Separase and APC were found to be necessary to disengage sister centrioles of centrosomes purified from HeLa cell that were added to *Xenopus* egg extract (Tsou and Stearns, 2006). Direct involvement of cohesin in centriole engagement was demonstrated in human culture cells (Schöckel et al., 2011). Mutations in cohesin and separase induce premature centriole disengagement in *C. elegans* spermatocyte during meiosis II (Schvarzstein et al., 2013). All of these studies established a concept that common molecules regulate the separation of sister centrioles and chromatids. Although the APC-separase pathway plays a decisive role in chromatid separation, the pathway may be dispensable in centriole disengagement. This is consistent with the fact that in *C. elegans* embryo, mechanical force can break engagement of the centrioles (Cabral et al., 2013). Therefore, the role of the APC-separase pathway in the regulation of centrosome number *in vivo* remains unclear.

Spermatogenesis in *C. elegans* occurs via 2 consecutive cell divisions of a spermatocyte to form 4 spermatids and 1 residual body (L’Hernault, 2006; Peters et al., 2010; Ward et al., 1981). The division of primary spermatocytes is similar to normal equatorial cell division, in which the contractile ring constricts after chromosome segregation, except that the constriction is not completed. Secondary spermatocytes remain connected with a cytoplasmic bridge, and the ring regresses later. The second cell division is unique because each secondary spermatocyte (connected with its sister spermatocyte) forms 2 buds (spermatids) and segregates a haploid set of the genome into each spermatid, which will then become a sperm. The 2 mother cells retract the constriction between each other and become a residual body, which do not contain genomic DNA, but show presence of actin, tubulin, ribosomes, and ER/Golgi (Winter et al., 2017). The 2 pairs of centrioles function as spindle poles in meiosis I and disengage in anaphase I. The disengaged centrioles duplicate before meiosis II to produce 4 pairs of centrioles, each distributed to the sperm after budding division. These centrioles do not undergo disengagement in anaphase II, due to the lack of separase at the centrosomes (Schvarzstein et al., 2013). The engaged pair of centrioles in sperm is disengaged after fertilization and duplicates to form a bipolar spindle in one-cell stage of the zygote (Pelletier et al., 2006).

In this study, we investigated the role of APC in the regulation of centrosome number during spermatogenesis in *C. elegans*. Involvement of APC in chromatid separation in *C. elegans* has been demonstrated for meiosis in oocyte and during spermatogenesis (Golden et al., 2000). Worms with mutation in *emb-27* or *emb-30* genes, which encode the subunits of APC, display defects in chromosome segregation and produce anucleated sperms (Sadler and Shakes, 2000). *C. elegans* APC might also regulate the engagement of sister centrioles during spermatogenesis. A previous study reported that zygotes obtained by mating *emb-27* mutant males with normal hermaphrodites showed cytokinesis with multiple furrows (Sadler and Shakes, 2000), which suggests that there is an excess number of spindle poles, and further, an excess number of centrosomes in the zygote with the paternal *emb-27* mutation. However, the centrosomes were not enumerated, and it remained unclear whether these zygotes possessed supernumerary centrosomes. Further, it is not known whether the excess number of centrosomes were brought into the zygote by the sperm, which could suggest a defect in centriole engagement in the *emb-27^APC6^* mutant. However, based on current view, APC activates separase to cleave cohesin and thus promote disengagement of the sister centrioles. This suggests that APC mutants should have a defect in centriole separation and thus centriole duplication. Therefore, APC mutation would be expected to result in decreased centrosome number, instead of the above-mentioned phenotype of an increase in centrosome number.

Thus, in this study, we attempted to solve two specific questions: (i) Does *emb-27* mutation result in an increased number of centrosomes in sperms and subsequently in *C. elegans* zygotes? (ii) How are the excess centrioles incorporated in the *emb-27* sperms during spermatogenesis? To address these questions, we conducted live cell imaging of spermatogenesis in *C. elegans*.

## Results

### Excess number of centrosomes in paternal *emb-27* mutant embryos

To examine the number of centrosomes in paternal *emb-27* (*g48ts*) mutant zygotes, we mated *emb-27* males grown at restrictive temperature with feminized hermaphrodites carrying a GFP-tagged γ-tubulin and observed the zygotes. γ-tubulin is one of the pericentriolar components of the centrosome (Bobinnec et al., 2000). As expected, we observed 3 or 4 centrosomes in the paternal *emb-27* mutant zygotes, with the frequency of cells with more than 2 centrosomes being almost 70% (Fig. 1A, B). This frequency of ∽70% was unexpected because the previous study by Sadler and Shakes (2000) described multiple furrows only in one-third of the embryos. Thus, excess number of centrosomes does not necessarily cause multiple furrows. We have further investigated the relationship between the excess number of centrosomes and the number of cytokinesis furrows in another study (Kondo and Kimura, 2018 preprint).

**Figure 1.**
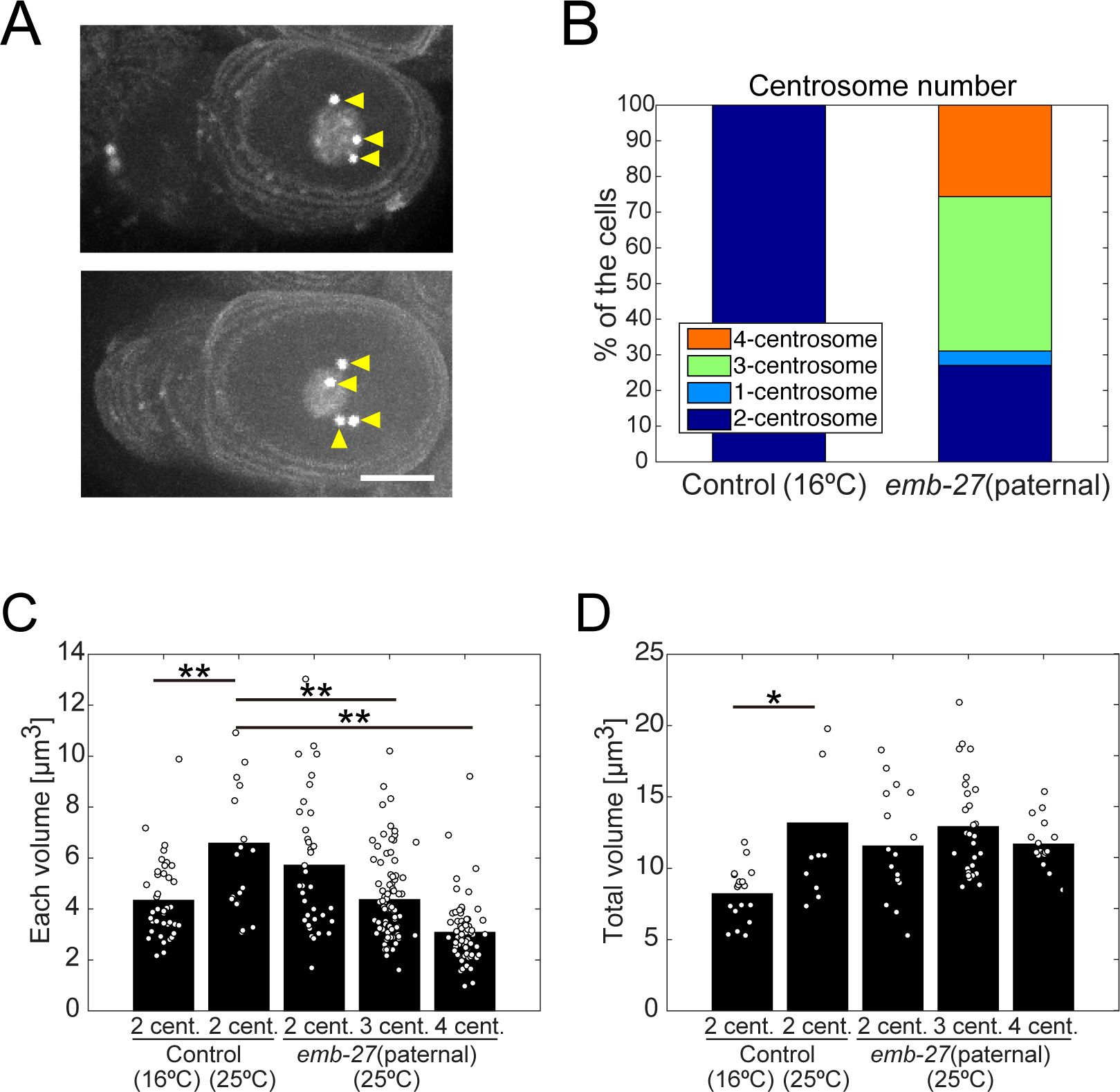
Abnormal number of centrosomes in *emb-27* paternal mutant embryos and spermatids. (A) Representative images of the 1-cell stage embryos of *emb-27* paternal mutant expressing GFP-tagged γ-tubulin (centrosome), PH^PLCδ1^ (cell membrane), and histone H2B (nucleus) *in utero*. Arrowheads indicate centrosomes in embryos with 3 (upper) and 4 centrosomes (lower). Bar, 10 μm. (B) The incidence of 1-cell embryos with the designated number of centrosomes. *n* = 22 (control) and 74 (*emb-27*). (C) Volumes of each centrosome in 1-cell embryos at metaphase. \*\**p* < 0.01 (Wilcoxon signed-rank test). (D) Total sum of centrosome volumes in 1-cell embryos at metaphase. \**p* < 0.05 (Wilcoxon signed-rank test). *n* = 40 (control, 16°C), 20 (control, 25°C), 36 (*emb-27*, 2-centrosome), 87 (*emb-27*, 3-centrosome), and 72 (*emb-27*, 4-centrosome).

ZYG-1 is a protein kinase, whose substrates include SAS-6, a key molecule for centriole biogenesis (Kitagawa et al., 2009). In the *C. elegans* zygote, paternal mutations of *zyg-1^Plk4^* gene (class II mutations) have been reported to cause supernumerary centrosomes, with the number varying from 1 to 10 (Peters et al., 2010). In contrast, the number of centrosomes in the *emb-27* mutation was 4 or less (Fig. 1B), indicating that the mechanism regulating centrosome number is different between *zyg-1* and *emb-27*.

### The total volume of centrosome is conserved

Since paternal *emb-27* mutant zygotes frequently showed excess number of centrosomes (Fig. 1), we tested the limited pool hypothesis for the regulation of centrosome size (Decker et al., 2011). According to this hypothesis, the total volume of centrosomes in an embryo is conserved during the early stages of embryogenesis regardless of the number of centrosomes in each cell. Thus, when the centrosome number decreases or increases due to *zyg-1* mutations, the sizes of individual centrosomes change accordingly and maintain the total volume (Decker et al., 2011). However, being embryonic lethal mutations directly involved in centrosome biogenesis, it cannot be excluded that the *zyg-1* mutations directly affected the centrosome size independent of the centrosome number. We tested the limited pool hypothesis using the paternal *emb-27* mutant. Unlike in the *zyg-1* mutants, the centrosomes duplicated and cells divided in the paternal *emb-27* embryos (Fig. S1), and thus the centrosome function was normal. We measured the diameter of the centrosomes in paternal *emb-27* embryos and calculated their spherical volume from the diameter. The volume of individual centrosomes in cells with 3 and 4 centrosomes was smaller than that in cells with 2 centrosomes (Fig. 1C, *p* < 0.01). Importantly, the total volume of centrosomes was almost constant regardless of the number of centrosomes in each cell (Fig. 1D, *p* > 0.05), providing additional support for the limited pool hypothesis.

Furthermore, we found that the total volume of the centrosomes depends on the culture temperature (Fig. 1D). The total volume was significantly decreased at 16°C than that at 25°C in control cells. Our observation indicates that the cell volume is not the sole determinant of the total volume of the centrosomes. Additionally, because the embryos developed to adulthood at both temperatures, embryonic development seems to be robust against centrosome volume fluctuations.

### The *emb-27* mutant sperm carries excess number of centrosomes

To investigate the cause of paternal *emb-27* embryos possessing excess number of centrosomes, we examined whether these were in excess also in the *emb-27* mutant sperms. We visualized the SAS-4 protein, a component of the centriole (Kirkham et al., 2003; Leidel and Gönczy, 2003), in sperms by immunostaining (Fig. 2A). In control sperms of *emb-27* males grown at a permissive temperature (16ºC), one large focus of SAS-4 signal was observed adjacent to the nucleus, as reported previously for normal sperms (Delattre et al., 2004; Kirkham et al., 2003; Leidel and Gönczy, 2003) (Fig. 2A, “16ºC”). This large focus should contain a pair of centrioles, which could mature to become 2 centrosomes in the embryo. In contrast, the *emb-27* mutant sperms grown at a restrictive temperature (25ºC) displayed multiple foci of SAS-4 (Fig. 2A, “25ºC”). Notably, most of the *emb-27* sperms did not possess a nucleus, which is consistent with the previous study (Sadler and Shakes, 2000). Further, in *emb-27* sperms, a pair of 2 small foci was often positioned in close proximity to each other. The small foci of SAS-4 signal in *emb-27* sperms were apparently smaller than those in the control, each of which corresponds to one centrosome (i.e., 2 centrioles). We suspect that each of these small foci in *emb-27* sperms corresponded to one centriole, rather than a pair of centrioles. This was further demonstrated by tracking the foci in the zygotes after fertilization. In a control zygote, 1 large focus (containing 2 centrioles) duplicates after fertilization and becomes 2 mature centrosome in zygotes (Schvarzstein et al., 2013). In contrast, 1 small focus (suspected to possess 1 centriole) from an *emb-27* sperm generates 1 mature centrosome in zygote (Fig. 2B). If we assume a large focus contains 2 centrioles and a small focus contains 1 centriole, the predicted number of centrioles in an *emb-27* sperm ranged from 2 to 4, and the frequency of appearance (Fig. 2A) was similar to that of mature centrosomes in *emb-27* zygote (Fig. 1B). These results demonstrated that an excess number of centrioles (i.e., > 2) are present in the majority of the *emb-27* sperms, and brought into the zygotes upon fertilization.

**Figure 2.**
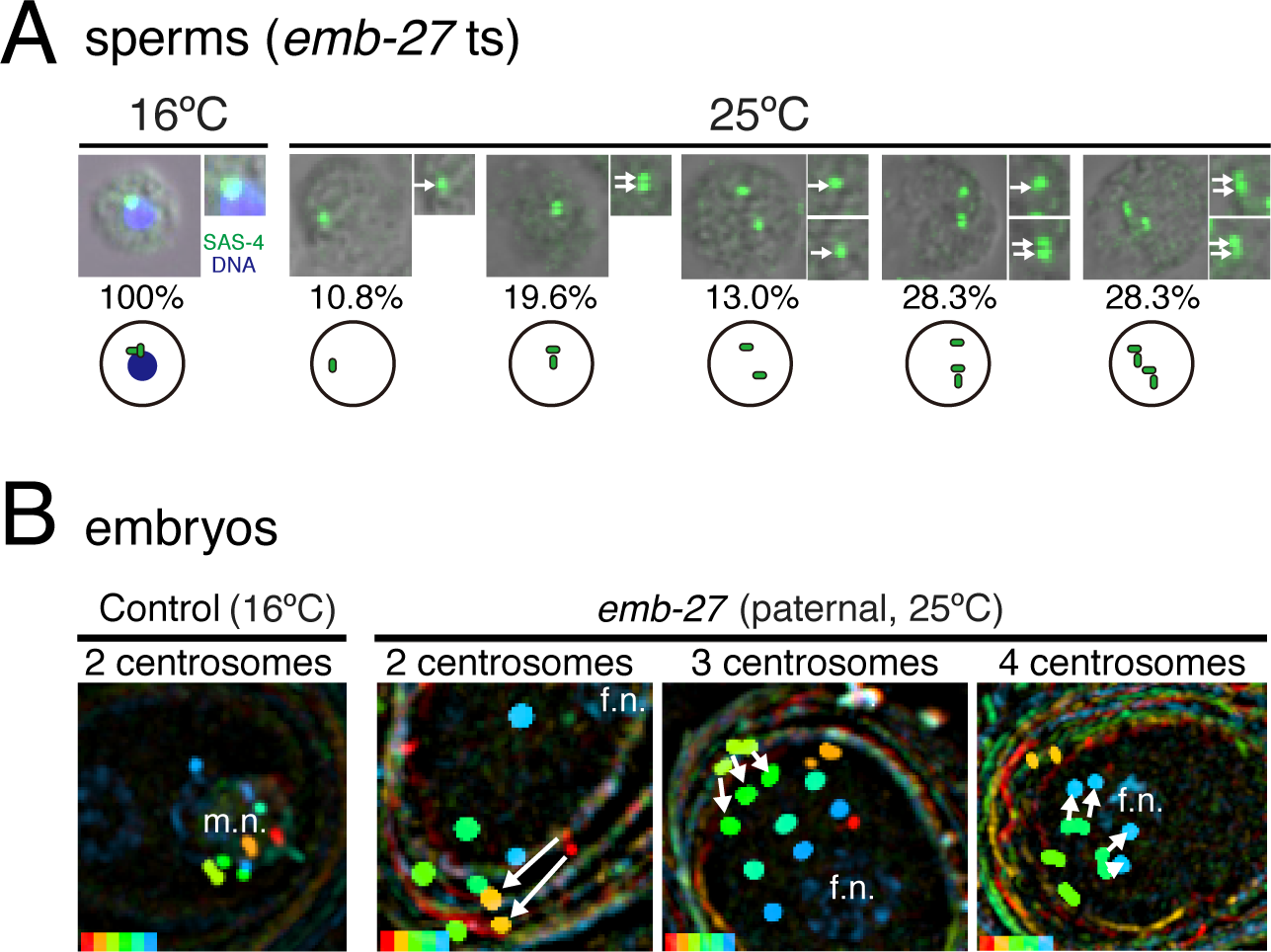
Abnormal number of centrosomes in *emb-27* paternal mutant spermatids. (A) Immunostaining of SAS-4 (green) in spermatids isolated from *emb-27*ts mutant males cultured at 16ºC (control) or 25ºC. DNA counterstained with DAPI (blue) and Nomarski differential phase contrast was also imaged. Arrows indicate SAS-4 foci. The percentages show the proportion of each SAS-4 focus pattern (*n* = 29 at non-restrictive temperature, and 46 at restrictive temperature). Cartoons show the putative centrioles (green). (B) Maturation process of centrosomes in 1-cell embryos. Each panel merges fluorescent signals of the centrosomes, nucleus, and cell membranes into one by showing the signals from different time points with different colors. Arrows indicate that adjacent GFP-tagged γ-tubulin spots are moving away from each other. m.n., male pronucleus; p.n., female pronucleus. In total, 22 and 67 individuals were observed for control and *emb-27* paternal mutants (*n* = 19 for 2-centrosome, 32 for 3-centrosome, and 16 for 4-centrosome), respectively, and similar results were obtained.

### The number of buds (spermatids) was reduced in *emb-27* spermatocyte

To investigate the mechanism underlying the excess centrioles in *emb-27* sperms, we performed live cell imaging during spermatogenesis (Figs. 3A, 3B, 4A, Movies 1, 2). In *emb-27^APC6^* mutant spermatocytes, chromosome segregation was impaired for both the first (i.e., equatorial) and second (i.e., budding) cell divisions of the spermatocyte (Fig. 3C, Movie 3). This was consistent with a previous observation of fixed spermatocytes in *mat* (<u>m</u>etaphase to <u>a</u>naphase <u>t</u>ransition-defective) mutants (Golden et al., 2000), and was expected because the APC is required for chromosome segregation (Uhlmann et al., 1999). The cytokinesis of the first division was also impaired in *emb-27^APC6^* (Fig. 3C, Movie 3), which is consistent with the idea that cytokinesis is allowed only after chromosomal segregation (Mendoza and Barral, 2008).

**Figure 3.**
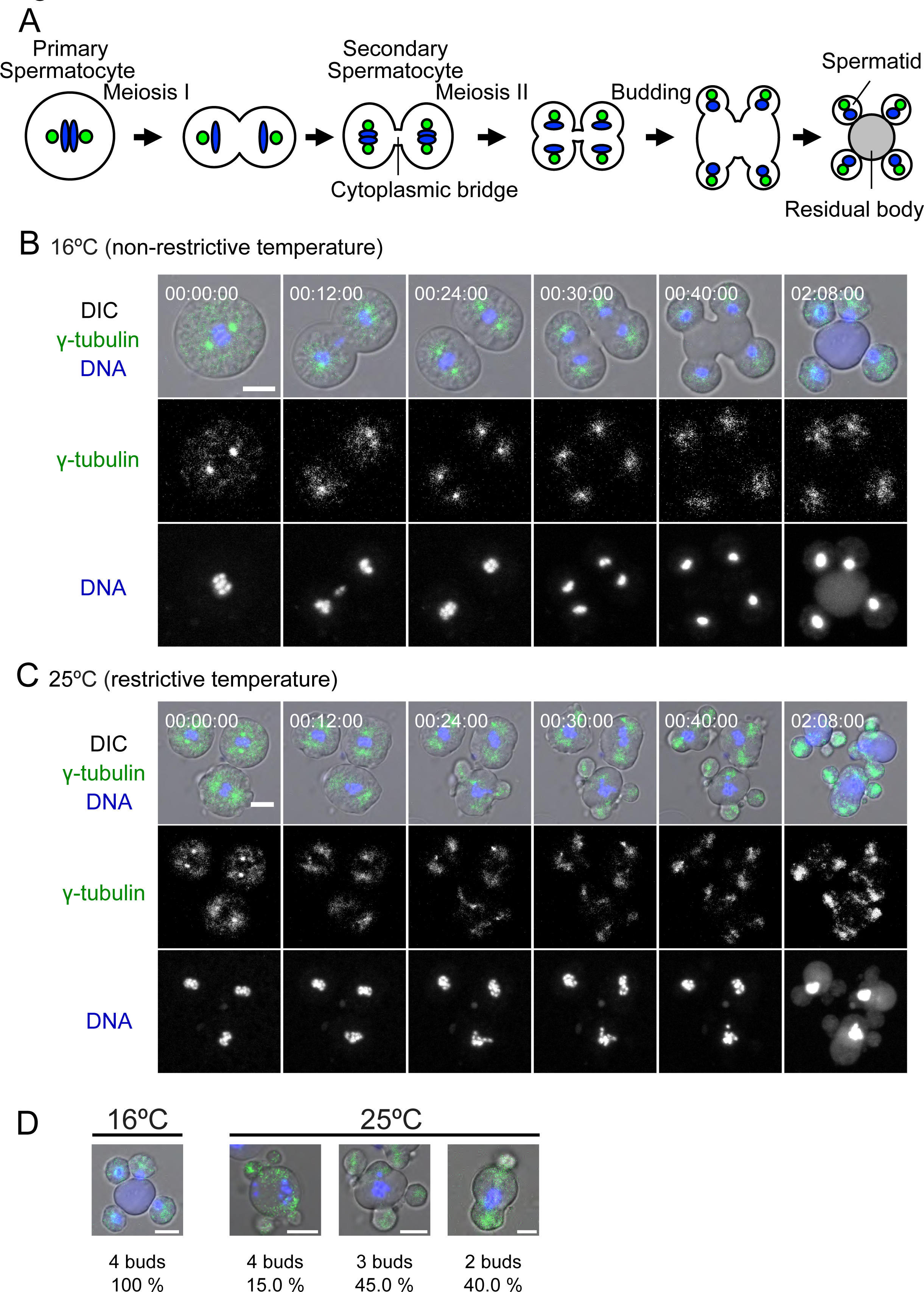
Spermatogenesis of *emb-27* paternal mutant. (A) Summary of spermatogenesis in *C. elegans*. In meiosis I, centrosomes (green) are located in the vicinity of chromatids (blue) and form a spindle in a primary spermatocyte (4N). Next, it generates two secondary spermatocytes (2N) that are linked by a cytoplasmic bridge after incomplete cytokinesis. Note that complete cytokinesis is also observed in this process (Ward et al., 1981). The secondary spermatocytes immediately initiate meiosis II to produce spermatids (N) and a residual body (grey) by budding. (B and C) Representative time-lapse images of a spermatocyte during spermatogenesis. The cells expressing GFP-tagged γ-tubulin were isolated from male CAL1151 (*emb27*ts) cultured at non-restrictive temperature (B) or restrictive temperature (C). DNA counterstained with Hoechst 33342. Bar, 5 μm. The elapsed time after the onset of imaging is indicated in hour:min:sec. In total, 9 and 22 individuals were observed at non-restrictive and restrictive temperatures, respectively, and similar results were obtained. (D) The number of buds per spermatocyte; 9 and 20 spermatocytes from *emb-27* mutant males at non-restrictive and restrictive temperature, respectively, were observed and the frequency of obtaining the indicated number of buds was calculated. Representative images are shown. Bar, 5 μm.

In contrast, cytokinesis of the second division (i.e., budding) was not impaired in *emb-27* spermatocytes (Fig. 3C, Movie 3). Although chromosomes did not segregate and remained in the mother cell (i.e., residual body), budding itself was not impaired, leading to formation of buds (spermatids) without chromosomes. The formation of sperms lacking chromosomes is consistent with the report of anucleated sperms in *emb-27* mutants (Sadler and Shakes, 2000). All the buds formed in *emb-27* mutant contained the SAS-4 signal, indicating that the centrosome, unlike the chromosome, is required for budding.

Importantly, we noticed that the number of buds was less than 4 in most cases (Fig. 3C, 3D). This observation may explain why *emb-27* sperms possess excess centrioles. In wild type, 4 centrosomes (8 centrioles) are segregated into 4 buds (sperms), resulting in each sperm containing 2 centrioles. During *emb-27* spermatogenesis, however, 8 centrioles are segregated into less than 4 buds, such that each resulting sperm contains 2 or more centrioles. The average number of centrioles per *emb-27* sperm was ∽3.1 (Fig. 2A), and the average number of buds in *emb-27* spermatocyte was ∽2.75 (Fig. 3D). This calculation implied that the total number of centrioles in *emb-27* spermatocyte is roughly 8 and is not decreased compared with wild type. Thus, the *emb-27* mutants do not have a defect in centriole disengagement, which should lead to impaired centriole duplication, during spermatogenesis.

### Centrosome separation defect in the secondary spermatocyte of *emb-27* mutant

To further investigate the reduced number of buds in *emb-27* mutants, we performed time-lapse imaging. Our observations showed that, in the *emb-27* mutant spermatocytes, centrosomes did not separate after meiosis I, and remained as 2 clusters at the onset of budding (00:12:00 in Fig. 3C), while the control spermatocytes contained 4 clusters of the centrosomes at the onset of budding (00:24:00 in Fig. 3B). As bud formation appears to be led by the centrosomes, the less number of the centrosome clusters is likely the cause of less number of buds in *emb-27* mutants.

In *C. elegans* embryo, a pair of sister centrosomes is separated toward opposite poles of the nucleus by sliding on the nucleus surface (Gönczy et al., 1999). In the *emb-27* spermatocyte, due to the defect in chromosome segregation (Fig. 3C), the number of nucleus remains 1 after meiosis I, and thus there are only 2 poles of 1 nucleus for 4 centrosomes. This implied that each of the 2 pairs of sister centrosomes cannot separate but remains clustered at one of the poles of the nucleus, which may get resolved during the budding stochastically (Fig. 3C). The stochastic resolution of the 2 sister centrosomes explains the formation of odd number of buds per spermatocyte (Fig. 3D).

### Nocodazole treatment at meiosis II inhibited centrosome separation and reduced the number of buds

To test our hypothesis that the defect in centrosome separation is the cause of decreased number of buds in *emb-27*, we impaired centrosome separation by using nocodazole, an inhibitor of microtubule polymerization. Upon application of nocodazole (Fig. 4Ba, Movie 4), centrosome separation was impaired in primary spermatocytes and 2 clusters of the centrosomes were observed throughout the imaging, as expected. This was in contrast to control spermatocytes or cytochalasin D (an inhibitor of actin polymerization)-treated spermatocytes, in which we observed 4 clusters of centrosomes at similar stages (Fig. 4A, 4Bc, Movies 1, 6). Unfortunately, we could not evaluate the relationship between centrosome separation and bud number in this experimental condition because the treatment with nocodazole of primary spermatocyte impaired bud formation completely (Fig. 4Ba).

**Figure 4.**
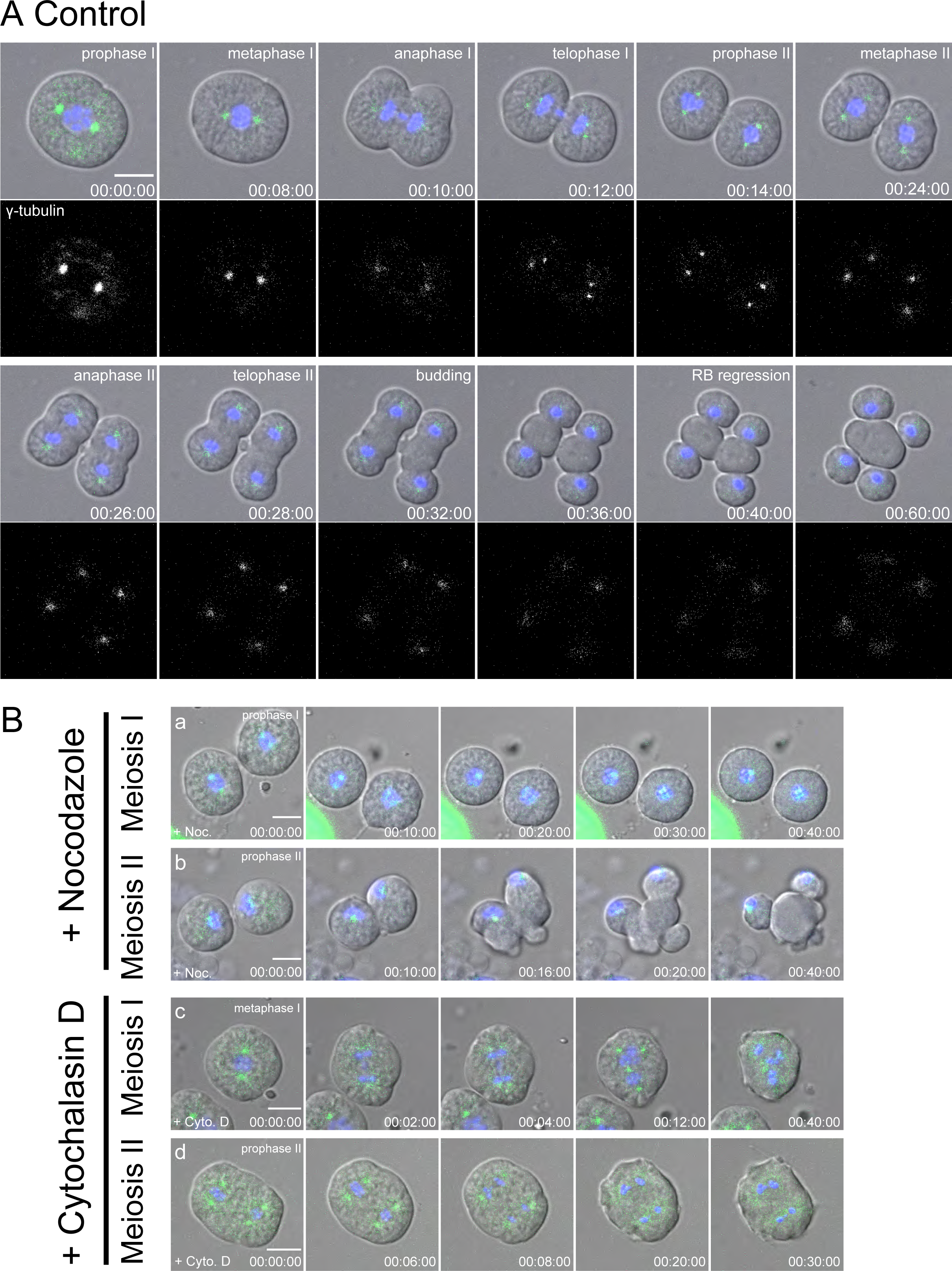
Live imaging of control spermatocyte treated with drugs. (A) DMSO (1%)-treated spermatocytes from control strain, CAL1121. DNA was counterstained with Hoechst 33342. Bar, 5 μm. In total, 4 individuals were observed and similar results were obtained. (B) Inhibition of actin and microtubule cytoskeleton by drugs. The cells were treated with the designated drugs several minutes before the onset of imaging, which was around prophase I (a), metaphase I (c), or prophase II (b and d). DNA was counterstained with Hoechst 33342. Bar, 5 μm. In total, 3 individuals were observed at each stage and similar results were obtained. The elapsed time after the onset of imaging is indicated in hour:min:sec.

Next, we focused on nocodazole treatment of secondary spermatocytes (Fig. 4Bb, Movie 5). In this condition, Meiosis I had completed and 2 nuclei were detected. Due to nocodazole treatment, the centrosomes failed to separate into 2 poles of the nuclei and remained as 2 clusters, resulting in the formation of only 2 buds (Fig. 4Bb, n = 3/3). This was in contrast to control spermatocytes or cytochalasin D-treated spermatocytes, in which we observed 4 clusters of centrosomes at similar stages (Fig. 4A, 4Bd, Movies 1, 7). This observation indicated that the number of buds corresponds to the number of centrosome clusters, and supports our model that the defect in centrosome separation in *emb-27* is the cause of reduced bud number in the spermatocyte and excess centrosome number in sperms.

### Budding (the second division) does not require the second chromosome segregation, but redundantly requires the first chromosome segregation and a microtubule-dependent process at meiosis II

In general, the inhibition of microtubule function by nocodazole impairs chromosome segregation, which is often required for the completion of cytokinesis. In contrast, we observed that, although the second chromosome segregation was impaired in secondary spermatocytes treated with nocodazole, the budding cytokinesis occurred in the cell (Fig. 4Bb). This observation indicated that chromosome segregation is not a prerequisite for budding cytokinesis in *C. elegans* spermatocyte. This supports our earlier observations with *emb-27* spermatocyte, where budding was not impaired even though chromosome segregation was defective (Fig. 3C).

However, if chromosome segregation is not required for budding, then the question remained as to why budding was impaired in nocodazole treatment at an earlier stage (Fig. 4Ba). In this earlier stage, chromosome segregation was impaired for both meiosis I and II, similar to that seen in *emb-27* spermatocytes, but budding was impaired, unlike in *emb-27* spermatocytes. Thus, microtubules may have an independent role in budding cytokinesis in addition to the role in chromosome segregation.

To further investigate the role of microtubules in the budding division, independent from chromosome segregation, we treated secondary spermatocytes of *emb-27* mutant with nocodazole. Interestingly, unlike in the controls (Fig. 3C and Fig. 4Bb), budding in *emb-27* spermatocytes was sensitive to the nocodazole treatment at the secondary spermatocyte stage (Fig. 5). This result suggests the presence of two processes required redundantly for budding: (i) nocodazole- and *emb-27*-sensitive process at meiosis I, and (ii) nocodazole-sensitive but *emb-27*-insensitive process at meiosis II. For the former, the only process we can think of is chromosome segregation at meiosis I. The latter process is unclear at the moment (see Discussion), but indicates a novel microtubule-dependent process specific for budding cytokinesis.

**Figure 5.**
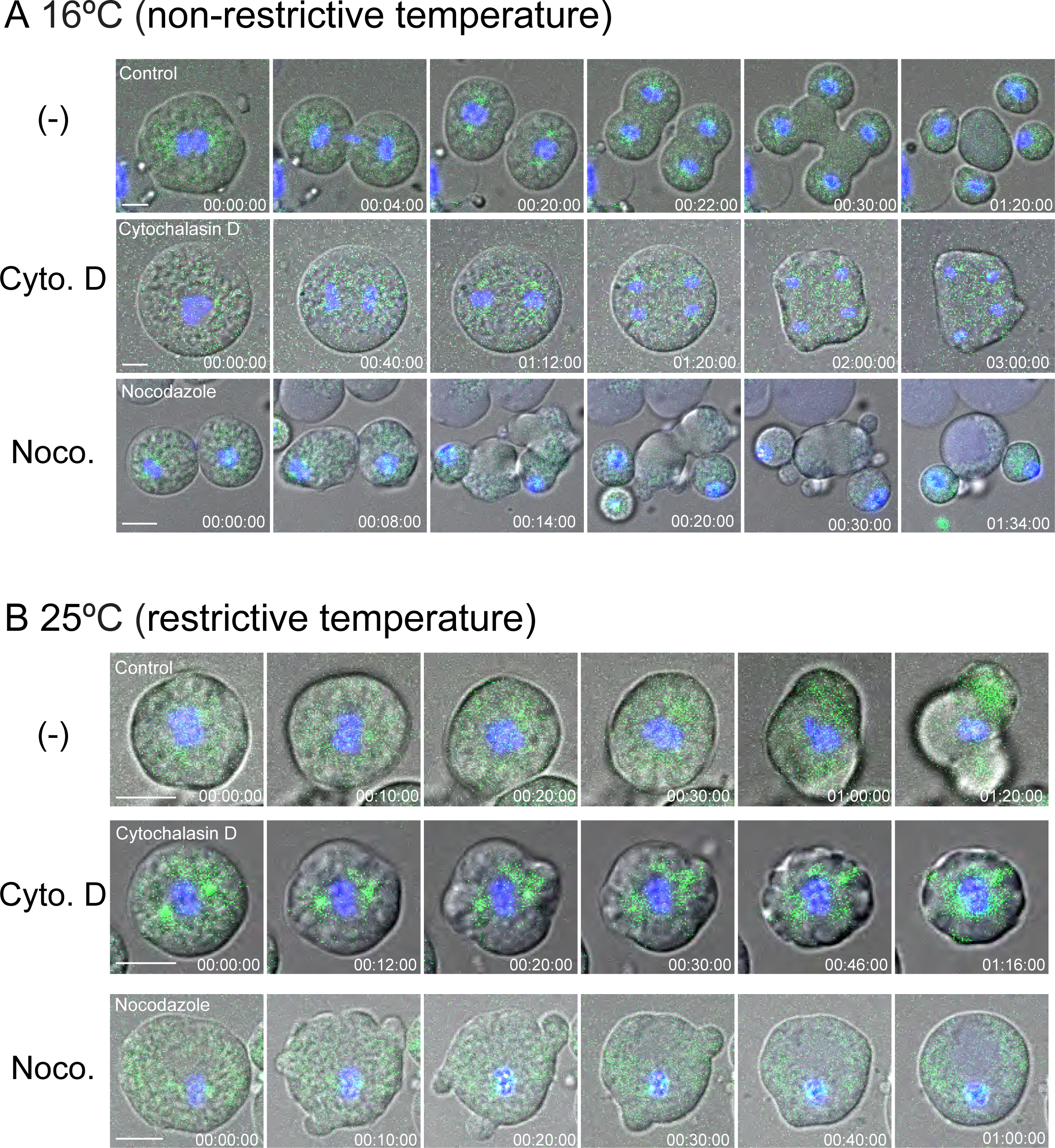
Live imaging of *emb-27* spermatocyte treated with drugs at meiosis II. (A and B) Spermatocytes from CAL1151 (*emb-27ts*; GFP::γ-tubulin [shown in green]) at non-restrictive temperature (A) or restrictive temperature (B). DNA was counterstained with Hoechst 33342 (blue); Bar, 5 μm. Cells were treated with the designated drugs several minutes before the onset of imaging, which was around prophase II. The experiment was repeated as follows: *n* = 5 for control, 4 for cytochalasin D (‘Cyto. D’) treatment, and 4 for nocodazole (‘Noco’) treatment at non-restrictive temperature, and *n* = 5, 7, and 9 for control, cytochalasin D, and nocodazole treatment, respectively, at restrictive temperature. Similar results were obtained.

## Discussion

In this study, we demonstrated that > 70% of *C. elegans* zygotes with paternal *emb-27* mutation had an anomalous number of centrosomes (Fig. 1), which was caused by an excess number of centrosomes incorporated into *emb-27* mutant sperms (Fig. 2). Based on our observations, we propose the following (Fig. 6): (i) an *emb-27* mutation, which can affect APC function, causes chromosome segregation defect in meiosis I and II, with the nucleus remaining one throughout spermatogenesis. (ii) In a normal spermatocyte, the centrosomes separate such that 4 centrosomes are positioned at 4 poles in total of 2 nuclei. However, as there is only one nucleus in the *emb-27* spermatocyte, the 4 centrosomes are positioned at 2 poles as 2 clusters. (iii) The number of buds is defined by the number of the centrosome clusters. As the *emb-27* spermatocyte has 2 clusters, 2 buds tend to appear. Our observation further suggested that 2 centrosomes in a cluster tend to separate during the budding to produce 3 or 4 buds per spermatocyte in a stochastic manner.

**Figure 6.**
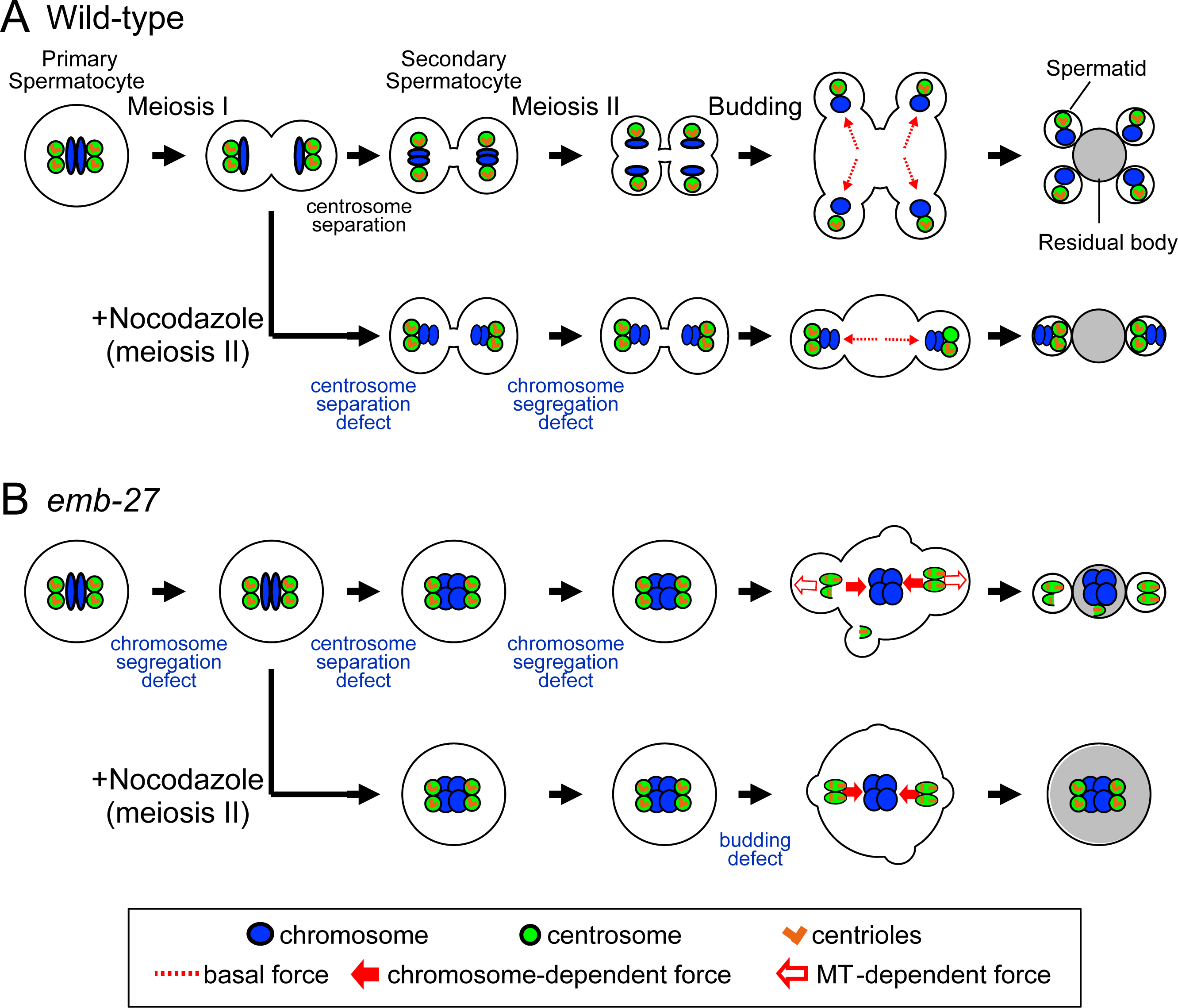
Proposed model for the mechanisms of generating sperms with excess centrosomes in *emb-27* mutant. Chromosome segregation defect caused by the mutation in *emb-27*^APC6^ also induces centrosome separation defect. Reduced number of centrosome clusters results in reduced number of buds. Based on the budding defect in *emb-27* spermatocyte treated with nocodazole at meiosis II, we propose the existence of 3 kinds of forces. The ‘basal’ force (red, dotted arrows) moves centrosomes (green circles) and chromosomes (blue) to the future buds in wild-type (A). Microtubule-dependent forces (red-outlined arrows) is required to move the centrosomes to the bud in *emb-27* mutant but not in wild-type. Chromosome-dependent forces (red-filled arrows) are likely pulling the centrosomes toward the center when the chromosome segregation is impaired in the *emb-27* spermatocyte. In the *emb-27* mutant, the engagement between centrioles (orange bars) is also weak, as suggested by the inclusion of an odd number of centrioles in the sperm.

The stochastic separation of the centrosome clusters may also occur between the sister centrioles. The odd number of 3 centrioles was predominantly found in an *emb-27* sperm (Figs. 1B, 2A), indicating that sister centrioles are separated at some point during the spermatogenesis in *emb-27* mutant. In fact, while two centrioles appeared as one large focus in the control sperm, we often observed a pair of small and adjacent foci of centrosomes in the *emb-27* sperms (Fig. 2A), suggesting that the link between a pair of centrioles was weak in the *emb-27* sperms. A premature disengagement of the sister centrioles has been reported in *C. elegans* sperms upon loss-of-function mutation of cohesin (*rec-8*(*ok978*)) and in a gain-of-function mutation of separase (*sep-1*(*e2406*)) (Schvarzstein et al., 2013). This observation is consistent with the idea that cohesin is involved not only in sister chromatid cohesion but also in sister centriole engagement, which is resolved by the APC-separase pathway (Schöckel et al., 2011; Tsou and Stearns, 2006). Based on the similarity between the phenotype and a well-established linkage between APC-separase-cohesin, we initially speculated that the *emb-27*(*g48*) mutation promoted centriole disengagement through premature/ectopic cleavage of cohesin. However, the *emb-27*(*g48*) mutation is a loss-of-function mutation inhibiting cohesin cleavage, at least for sister chromatid cohesion. The underlying mechanism responsible for the observed pairs of small foci of centrosome signal in *emb-27* is likely different from that in the *rec-8* and *sep-1* mutants, because an extra centriole duplication occurred in these two mutants after the premature disengagement of centrioles (Schvarzstein et al., 2013), but not in the *emb-27* mutant as judged from the number of centrosomes in the zygotes (Fig. 1, 2). In the present study, we did not obtain any evidence for the involvement of *emb-27*^APC6^ in centriole engagement except for the microscopic observation of separated centrosomal foci in sperms (Fig. 2A). If the *emb-27* mutation affects centriole engagement, it should affect centriole duplication and the number of centriole should be abnormal, which was not the case. Thus, our study suggest that the APC-separase-cohesin pathway may be dispensable for the regulation of centriole engagement in *C. elegans* spermatogenesis, as indicated in other systems (Cabral et al., 2013; Oliveira and Nasmyth, 2013).

The present study suggested a novel role of microtubule in the second cell division (budding cytokinesis) of spermatocyte. The first cell division (furrowing cytokinesis) in spermatogenesis appeared to be very similar to other types of cell division such as embryonic cell division except that the cleavage was incomplete. This cell division was dependent on actin, microtubule, and APC function. In contrast, the second cell division (budding cytokinesis) was actin-dependent, but was not inhibited by microtubule drug or *emb-27* mutation. In the *emb-27* mutant, chromosomes remained in the mother cell (residual body), indicating that budding cytokinesis does not require chromosome segregation. Further, budding was observed when wild type secondary spermatocytes were treated with nocodazole (microtubule depolymerizing drug), indicating that microtubule elongation is not required for budding. In contrast, budding did not occur when *emb-27* secondary spermatocytes were treated with nocodazole. This indicates that two processes are required redundantly for the budding – one is a nocodazole- and *emb-27*-sensitive process at meiosis I, and the other is a nocodazole-sensitive but *emb-27*-insensitive process at meiosis II. Chromosome segregation at meiosis I should correspond to the former process. For the latter, we propose a microtubule-based force counteracting with a force pulling the centrosomes towards the chromosomes. In *emb-27* spermatocytes, we often observed an attraction between chromosomes and centrosomes (Movie 3). The centrosomes moving toward a growing bud were often pulled back toward the chromosomes in the mother cell, which caused retraction of the bud or fragmentation of the centrosomes (Fig. 3C, Movie 3). To overcome this attraction from the mother cell and move toward the periphery to form buds in *emb-27* spermatocytes, the centrosomes require a force in the direction of the buds. This force could be microtubule-dependent, as the centrosomes do not leave the chromosomes in nocodazole-treated *emb-27* spermatocytes (from meiosis II, Fig. 5B) or control spermatocytes from meiosis I (Fig. 4Ba). This microtubule-based force is indispensable for budding when chromosome segregation at meiosis I was successful (Fig. 4Bb).

In summary, we propose the existence of three kinds of forces to move the centrosomes to the bud (Fig. 6). The ‘basal’ force (red, dotted arrows in Fig. 6), possibly dependent on actin, moves centrosomes and chromosomes to the future buds in wild type. Based on our nocodazole treatment experiments, in the *emb-27* mutant spermatocyte, budding is driven by microtubule-dependent forces (red-outlined arrows in Fig. 6). Because the microtubule-dependent forces are dispensable in wild type, chromosome-dependent forces (red-filled arrows in Fig. 6) induced by the defect in chromosome segregation, are likely pulling the centrosomes toward the center in the *emb-27* spermatocyte.

These forces also explain the formation of 3 or 4 buds per spermatocyte and the incorporation of the odd number of centrioles into a sperm in *emb-27* mutants. In the secondary spermatocyte of *emb-27*, the separation of the two centrosomes to a pole of the nucleus, and the separation of the sister centrioles in a centrosome were observed, which were not observed in nocodazole treated spermatocytes. The two opposing forces acting on the centrosomes in *emb-27*—the nocodazole-sensitive force to move the centrosomes toward the sperm bud, and the other is to retain the centrosomes toward the residual body by the chromosomes there—may mechanically separate the centrosomes and the centrioles. It has been shown that mechanical force can break engagement of the centrioles in the *C. elegans* embryo (Cabral et al., 2013). A further improvement in the resolution in time and space to follow the behavior of centrosomes during spermatogenesis might provide hints to reveal the underlying mechanism.

## Materials and methods

### Worm strains and maintenance

The strains used in this study are listed in Table S1. Strains were maintained under standard conditions (Brenner, 1974). CAL0051 was obtained by crossing GG48 and DR466. CAL0182 was obtained by crossing OD58 and TH32, and then with BA17. CAL1151 was obtained by crossing CAL0051 and TH32, and CAL1121 was obtained by crossing DR466 and TH32. The generation of the desired genotype was checked by determining whether 100% of the progeny showed the desired fluorescence(s) or mutant phenotype(s). To obtain *emb-27* mutant males, temperature shift was conducted as follows: 10 gravid hermaphrodites of *emb-27*ts (CAL0051, or CAL1151) were placed on a fresh 60-mm plate and allowed to lay eggs at 16ºC (non-restrictive temperature). After 21-28 h, the adults were removed from the plate, and the plate was incubated for another 26-30 h at 16ºC. Next, the plate was transferred to 25ºC (restrictive temperature) and incubated for another 13-18 h before mating or observation of spermatocytes/sperms. In Fig. 2A, 3B and 5B, *emb-27* at ‘16°C (non-restrictive temperature)’ indicates that the *emb-27*ts males were consistently grown at 16°C as a control.

To obtain paternal *emb-27* mutant embryos (Figs. 1, 2B, and S1), 30 *emb-27* mutant males from the plate were moved to a new 35-mm plate with 10 young adult hermaphrodites of CAL0182 or CAL0041 for 13-18 h at 25ºC to induce mating, and the embryos cut out from the hermaphrodites were observed. CAL0182 (*fem-1*ts) from the L1/L2 stage was grown at 25ºC to prevent self-fertilization. The control embryos were obtained from adult hermaphrodites of CAL0182 or CAL0041 that were grown at 16°C until young adults to allow self-fertilization. The ‘control (25°C)’ in the figures indicates that the control hermaphrodites were incubated at 25°C for 13–18 h prior to observation to achieve conditions similar to those for the *emb-27* paternal mutant embryos. ‘Control (16°C)’ in the figures indicates that the control hermaphrodites were consistently incubated at 16°C.

### Immunofluorescence microscopy of male sperms (Fig. 2A)

Adult males of CAL0051 incubated prior to observation at 16ºC or temperature-shifted to 25ºC for 13-18 h were anesthetized with 2 mM (-)-tetramisole hydrochloride (L9756; Sigma-Aldrich, St Louis, MO, USA) in M9 solution (Brenner, 1974) for 3-5 min and transferred to a drop of sperm medium (50 mM HEPES titrated to pH 7.0 with NaOH, 50 mM NaCl, 25 mM KCI, 5 mM CaCl2, 1 mM MgSO4, and 1 mg/ml BSA) (Nelson and Ward, 1980). The worms were then dissected and gently punched by a proximal arm on a poly-L-lysine (P8920; Sigma-Aldrich)-coated glass slide (Multitest slide 8-well; MP Biomedicals LLC, Solon, OH, USA). The bared sperms were fixed with -20ºC methanol, covered with a coverslip, and made permeable by freezing on dry ice for more than 30 min followed by removing the coverslip. Indirect immunofluorescence was conducted according to a previously described method (Matsuura et al., 2016) with some modifications. After washing with PBS (137 mM NaCl, 2.7 mM KCl, 10 mM Na2HPO4, 2 mM KH2PO4) containing 0.05% Tween-20 (PBS-T) for 5 min, the gonads were treated with 1% BSA in PBS for 20 min and incubated with primary antibodies for SAS-4 (sc-98949; Santa Cruz Biotechnology, Santa Cruz, CA, USA) and α-tubulin (T6199; Sigma-Aldrich) diluted with 1% BSA in PBS for overnight at 4ºC in a humidity chamber. After subsequent washing with PBS-T three times for 5 min each, the gonads were treated with Alexa Fluor-conjugated secondary antibodies for anti-mouse IgG (A11004; Molecular Probes, Eugene, OR, USA) and anti-rabbit IgG (A11008; Molecular Probes) for 1 h at room temperature and rinsed with PBS-T three times for 5 min each. Next, the gonads in a drop of SlowFade Gold antifade reagent (S36936; Life Technologies, Carlsbad, CA, USA) containing 10 µg/ml Hoechst33342 (346-07951; Dojindo, Kumamoto, Japan) were covered with a coverslip. The specimens were observed using a laser scanning confocal microscope (FV-1200; Olympus, Tokyo, Japan) with an UPlanSApo 100× 1.4 NA objective (Olympus). Image processing was conducted using Fluoview (Olympus).

### Live cell imaging microscopy: Observation of spermatocytes in males (Fig. 3–5)

For observation of living spermatocytes, adult males of CAL1151 (*emb-27*ts, Figs. 3 and 5) or CAL1121 (control, grown at 22°C, Fig. 4) were dissected in a drop of the sperm medium containing 5 µg/ml Hoechst 33343 and sealed using VALAP. For inhibition of actin and tubulin, the dissection was performed in the sperm medium containing 5 µg/ml cytochalasin D (Sigma-Aldrich) or 5 µg/ml nocodazole (Sigma-Aldrich), respectively. Images were acquired every minute by using a laser scanning confocal microscope (FV-1200) with an UPlanSApo 100× 1.4 NA objective (Olympus). The fluorescence signals were detected using GaAsP detectors (Olympus).

### Live cell imaging microscopy: Observation of embryos

For observation of embryos *in utero* (Fig. 1A), anesthetized adult worms were placed on 2% agar pad and gently sealed with a coverslip (Kimura and Kimura, 2012). For observation of embryos *ex utero* (Fig. S1), embryos were obtained by cutting open gravid adult worms in M9 solution. The embryos were transferred to a drop of M9 solution on a glass slide (Multitest slide 8-well) and sealed using VALAP. The samples were observed using a spinning-disk confocal system (CSU-X1; Yokogawa Electric, Tokyo, Japan) mounted on a microscope (IX71; Olympus). Images were acquired every minute (for *in utero*) or at 5-min intervals (for *ex utero*) for the thickness of 30 μm with 2 μm z-intervals at 20 ms exposure by using a UPlanSApo 60× 1.3 NA objective (Olympus) equipped with an EM-CCD camera (iXon; Andor, Belfast, UK) controlled by Metamorph software (Molecular Devices, Sunnyvale, CA, USA).

### Image processing and analysis

For obtaining four-dimensional stack images shown in Fig. 2B, the 3D projected images were preprocessed using a Difference of Gaussian filter stacked with an ImageJ plugin Color Footprint Rainbow developed by Dr. Hiratsuka (JAIST, Ishikawa, Japan). For the measurement of centrosome size as shown in Fig. 1C–D, the obtained images were filtered with a Gaussian blur (σ = 1.0), and then a profile of each centrosome was determined using a 1-pixel wide line scan by using ImageJ. The profile was fit to the Gaussian function *y* = *a* + (*b* - *a*) × exp^-(*x* - *c*) × (*x* - *c*)/2 × *d* × *d*^ by using a custom-written ImageJ macro, and the value of *d* × 2 × (2 × ln2)^0.5^, the full width at the half maximum, was defined as a centrosome diameter (Greenan et al., 2010). The value was used to calculate a centrosome volume as a sphere.

### Statistical analyses

For the comparison of centrosome sizes from the two groups (Fig. 1C, 1D), we first tested the normality of the data by using one sample Kolmogorov-Smirnov test by using the MATLAB ‘kstest’ function. Since the distributions were not normal, we compared pairs of two groups by using Wilcoxon rank sum test (MATLAB ‘ranksum’ function).

## Acknowledgments

We thank Drs. Rieko Matsuura and Daiju Kitagawa (National Institute of Genetics) for technical advice on immunofluorescence of sperms, Ms. Yoko Kimura (National Institute of Genetics) for help with establishing the strains, Drs. Daiju Kitagawa, Yasushi Hiromi, Mitsuhiko Kurusu, Hiroaki Seino, Kenji Kimura and Yohei Kikuchi (National Institute of Genetics) for their comments on the manuscript, and all the members of our laboratory for discussion. Some worm strains used in this work were provided by the Caenorhabditis Genetics Center. T.K was a postdoctoral fellow of the National Institute of Genetics.

## Author contributions

T.K. and A.K. conceived the study, analyzed the data, and wrote the manuscript. T.K. performed all the experiments.

## Data availability

The raw data is available upon request.

## Supplementary Files

**Table S1.**
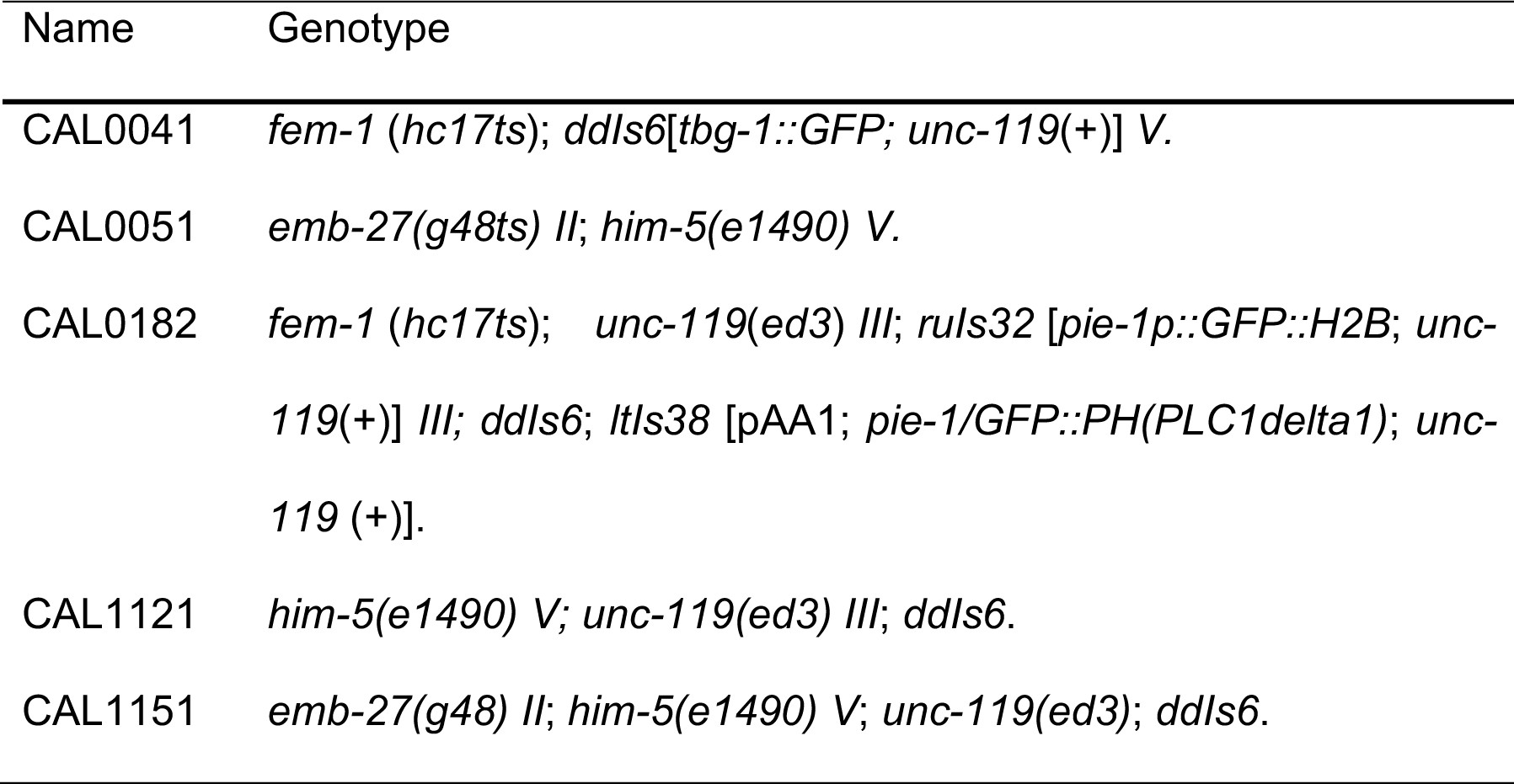
strains used in this study

**Figure S1.**
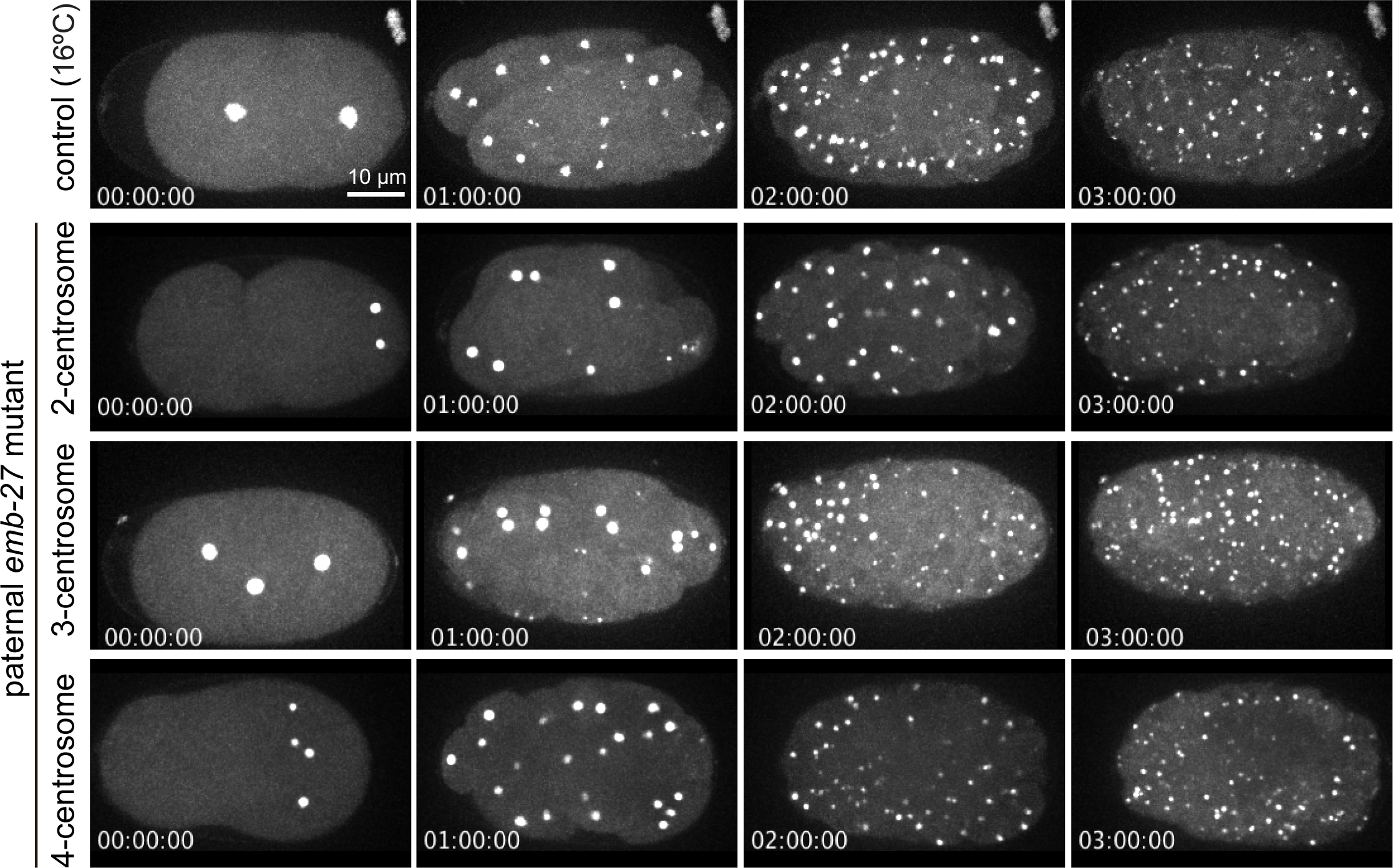
Long-term time-lapse images of emb-27 paternal mutant expressing GFP-tagged γ-tubulin. The ex utero embryos of CAL0041 strain were imaged every 5 min for 3 h. Each image is a maximum-intensity projection of 16 sections collected at 2 μm steps. In total, 11 and 10 individuals were observed for control and emb-27 paternal mutants (n = 3 for 2-centrosome, 4 for 3-centrosome, and 3 for 4-centrosome), respectively, and similar results were obtained.

**Movie 1. Time-lapse movie of a spermatocyte during spermatogenesis in the control strain without drugs (related to Fig. 4A)**

Time-lapse confocal microscopy images of DMSO (1%)-treated spermatocytes from control strain, CAL1121. TBG1::GFP signal is shown in green. DNA was counterstained with Hoechst 33342 (blue). Time is shown with hrs:min:sec. Bar, 5 μm.

**Movie 2. Time-lapse movie of a spermatocyte during spermatogenesis in *emb-27* at permissive temperature (related to Fig. 3B)**

Time-lapse confocal microscopy images of spermatocytes from CAL1151 (*emb27*ts) cultured at non-restrictive temperature. TBG1::GFP signal is shown in green. DNA was counterstained with Hoechst 33342 (blue). Time is shown with hrs:min:sec. Bar, 5 μm.

**Movie 3. Time-lapse movie of a spermatocyte during spermatogenesis in *emb-27* at restrictive temperature (related to Fig. 3C)**

Time-lapse confocal microscopy images of spermatocytes from CAL1151 (*emb27*ts) cultured at restrictive temperature. TBG1::GFP signal is shown in green. DNA was counterstained with Hoechst 33342 (blue). Time is shown with hrs:min:sec. Bar, 5 μm.

**Movie 4. Time-lapse movie of a spermatocyte during spermatogenesis (meiosis I) after nocodazole treatment (related to Fig. 4Ba)**

Time-lapse confocal microscopy images of nocodazole-treated spermatocytes during meiosis I from control strain, CAL1121. TBG1::GFP signal is shown in green. DNA was counterstained with Hoechst 33342 (blue). Time is shown with hrs:min:sec. Bar, 5 μm.

**Movie 5. Time-lapse movie of a spermatocyte during spermatogenesis (meiosis II) after nocodazole treatment (related to Fig. 4Bb)**

Time-lapse confocal microscopy images of nocodazole-treated spermatocytes during meiosis II from control strain, CAL1121. TBG1::GFP signal is shown in green. DNA was counterstained with Hoechst 33342 (blue). Time is shown with hrs:min:sec. Bar, 5 μm.

**Movie 6. Time-lapse movie of a spermatocyte during spermatogenesis (meiosis I) after cytochalasin D treatment (related to Fig. 4Bc)**

Time-lapse confocal microscopy images of cytochalasin D-treated spermatocytes during meiosis I from control strain, CAL1121. TBG1::GFP signal is shown in green. DNA was counterstained with Hoechst 33342 (blue). Time is shown with hrs:min:sec. Bar, 5 μm.

**Movie 7. Time-lapse movie of a spermatocyte during spermatogenesis (meiosis II) after cytochalasin D treatment (related to Fig. 4Bd)**

Time-lapse confocal microscopy images of cytochalasin D-treated spermatocytes during meiosis II from control strain, CAL1121. TBG1::GFP signal is shown in green. DNA was counterstained with Hoechst 33342 (blue). Time is shown with hrs:min:sec. Bar, 5 μm.

